# The Role and Mechanism of TEAD4 in Preimplantation Embryonic Development in Mice and Cattle

**DOI:** 10.1101/2023.07.13.548853

**Authors:** Xiaotong Wu, Yan Shi, Bingjie Hu, Panpan Zhao, Shuang Li, Lieying Xiao, Shaohua Wang, Kun Zhang

**Affiliations:** Laboratory of Mammalian Molecular Embryology, College of Animal Sciences, Zhejiang University, Hangzhou, Zhejiang 310058, China; Key Laboratory of Dairy Cow Genetic Improvement and Milk Quality Research of Zhejiang Province, College of Animal Sciences, Zhejiang University, Hangzhou, Zhejiang 310058, China

**Keywords:** Mouse, Bovine, Embryo, Base editing, Tead4, Trophectoderm

## Abstract

Tead4, a critical transcription factor expressed during preimplantation development, is essential for the expression of trophectoderm-specific genes in mice. However, the functional mechanism of *Tead4* in mouse preimplantation development and its conservation across mammals remain unclear. Here, we report that Tead4 is a crucial transcription factor necessary for blastocyst formation in mice. Disruption of *Tead4* through base editing results in developmental arrest at the morula stage. Additionally, RNA-seq analysis reveals dysregulation of 670 genes in *Tead4* knockout embryos. As anticipated, *Tead4* knockout leads to a decrease in trophectoderm genes *Cdx2* and *Gata3*. Intriguingly, we observed a reduction in Krt8, suggesting that Tead4 influences the integrity of the trophectoderm epithelium in mice. More importantly, we noted a dramatic decrease in nuclear Yap in outside cells for *Tead4*-deficient morula, indicating that Tead4 directly regulates Hippo signaling. In contrast, bovine embryos with *TEAD4* depletion could still develop to blastocysts with normal expression of *CDX2*, *GATA3*, and *SOX2*, albeit with a decrease in total cell number and ICM cell number. In conclusion, we propose that Tead4 regulates mouse blastocyst formation via Krt8 and Yap, both of which are critical regulators of mouse preimplantation development.

## Introduction

Upon fertilization, a totipotent zygote continuously divides to create a blastocyst, a process known as preimplantation embryonic development in mammals. Morphologically, the zygote undergoes cleavage, compaction, polarization, and cavitation to generate a blastocyst with three embryonic layers: the trophectoderm (TE), epiblast (EPI), and primitive endoderm (PE). The TE differentiates to form the placenta, the EPI gives rise to the fetus, and the PE develops into the yolk sac. Although the preimplantation development appears conserved across mammalian species, there are notable differences in regulation of key biological events within this period ^1–3^.

Tead4 is one of the four members of TEAD family transcription factors ^4^. *Tead4* mRNA begins to express faintly at the 2-cell stage, peaks at the 8-cell stage, and maintains its expression during the blastocyst stage. The knockout (KO) of *Tead4* through homologous recombination results in developmental arrest at the morula stage ^5–7^. Furthermore, the mRNA expression of TE marker genes *Cdx2*, *Gata3*, *Eomes*, *Fgfr2*, *Itga7*, and *Cdh3* decreased significantly, while the mRNA expression of ICM marker genes *Fgf4*, *Sox2*, and *Oct4* remained unaffected in *Tead4* KO embryos. It has been believed that Tead4 operates upstream of genes associated with TE specification and function ^8–11^. However, considering that these TE marker genes (*Cdx2*, *Gata3* etc.) are not essential for TE specification and blastocyst formation, it remains unclear how Tead4 leads to developmental arrest at morula stage in mice.

While the significance of Tead4 in regulating TE specification and differentiation in mice has been well established by KO studies, the role and regulatory mechanism in other mammalian species remains unclear. Recently, two studies showed that knocking down TEAD4 via RNA interference does not affect development to the blastocyst stage in cattle, however, they produced contrasting results regarding the role of TEAD4 in TE marker genes. Given the limitation of the RNAi approach in completely removing endogenous mRNA, the functional role of TEAD4 in lineage specification and blastocyst formation in bovine preimplantation development remains elusive.

In this study, we have created robust models of *Tead4* KO embryos to determine the mechanisms underlying the requirement of Tead4 in mouse embryos and whether TEAD4 is essential for blastocyst formation in cattle. Through RNA-seq and immunofluorescence analysis of the wildtype (WT) and KO embryos, we have unveiled the functional requirement of Tead4 in lineage segregation, integrity of TE epithelium, and Hippo signaling during blastocyst formation in mice. Interestingly, this is in stark contrast to the effects of bovine TEAD4, which is not required for blastocyst formation and lineage differentiation in cattle.

## Results

### Generation of Point Mutations in Mouse *Tead4*

To investigate mechanisms underlying the role of Tead4 in TE lineage specification in mouse preimplantation embryos, we first utilized cytosine base editor 3 (BE3), a CRISPR-based tool, to induce point mutations in mouse *Tead4*. BE3 could mediate the conversion of C-G base pairs to T-A base pairs (C to T, G to A) ^12^, thereby generating premature stop codons. We initially designed two guide RNAs (gRNAs) in mouse *Tead4* exon 1 and exon 7, anticipating that the stop codons would be produced by conversions of C4, C5 to T for gRNA1 and C6, C7 to T for gRNA2 at the target sites (Fig. 1A).

**Fig 1.**
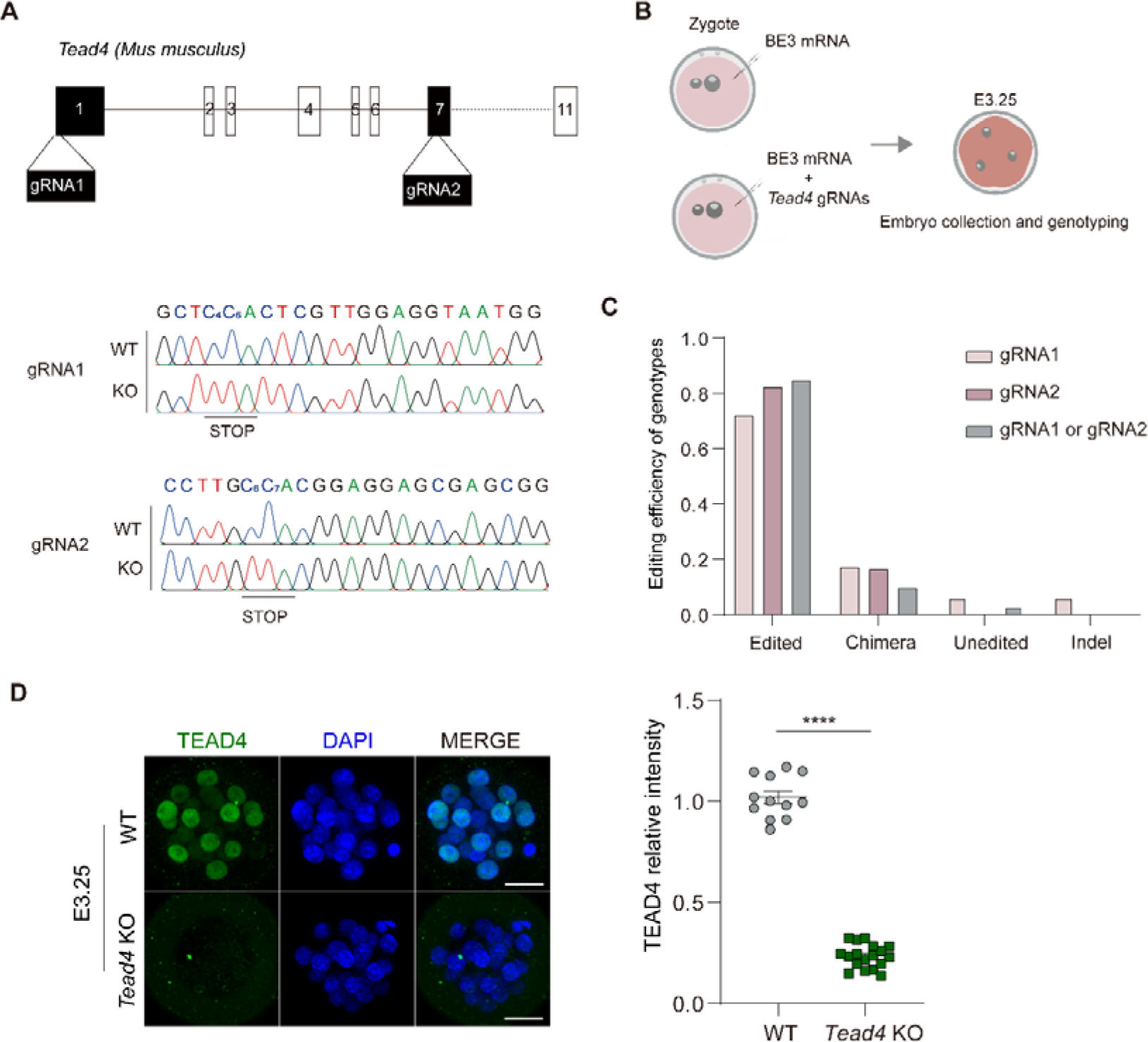
Generation of *Tead4* KO model in mouse embryo. (A) Two gRNAs were designed in mouse *Tead4* exon1 and exon7. C_4_, C_5_ in gRNA1 and C_6_, C_7_ in gRNA2 are potential target sites. (B) Scheme of genotyping in mouse *Tead4* knockout embryos. Zygotes were injected with BE3 mRNA or *Tead4* gRNAs and BE3 mRNA. Embryos were cultured to morula stage (E3.25) and subjected to genotyping. (C) Editing efficiency of different phenotypes in mouse *Tead4* KO embryos. Data are mean±s.e.m. (n=119). (D) Confocal images of TEAD4 in control and *Tead4* KO mouse embryos at morula stage. Scale bar, 25 μm. Quantification of the relative fluorescence intensity of TEAD4 in control and KO mouse embryos at morula stage. Data are mean±s.e.m. (n=30). ****: P<0.0001.

We subsequently microinjected a mixture of BE3 mRNA and gRNAs into mouse zygotes (Fig. 1B). To validate the efficiency of base editing, we collected mouse embryos at morula stage (E3.25) and conducted genotyping, revealing that 73.1% (76/104) of the embryos were correctly edited at two target sites for gRNA1, 84.5% (82/97) for gRNA2, and 89.9% (107/119) for gRNA1 or gRNA2 (Fig. 1C). The editing efficiency of other genotypes was significantly lower than these ones (Table. S1). Consistent with the change of genome, the nuclear intensity of Tead4 was significantly reduced in KO embryos (Fig. 1D). These results clearly demonstrate the robust editing efficiency of *Tead4* by BE3 in mouse preimplantation embryos.

### Tead4 is Essential for Mouse Blastocoel Formation

Given the high editing efficiency, we sought to explore the abnormalities in *Tead4* KO embryos. In WT embryos, we first detected TEAD4 with a weak signal at the four-cell stage, followed by an increase in immunostaining intensity from the eight-cell stage and maintained to the blastocyst stage (Fig. S1). We then tracked the preimplantation development from the two-cell stage to examine if there would be any morphological differences between *Tead4* WT and KO embryos. The *Tead4* KO embryos underwent compaction at the eight-cell stage and remained morphologically indistinguishable from the WT embryos until the morula stage. Then, the *Tead4* KO embryos arrested at the morula stage while the Tead4 WT embryos formed blastocyst (Fig. 2A). However, the cells of the *Tead4* KO embryos were found to proliferate at increased rates similar to those of the *Tead4* WT embryos until E3.25 (Fig. 2B). These results confirm that Tead4 plays an essential role in morula to blastocyst transition in mouse embryos.

**Fig 2.**
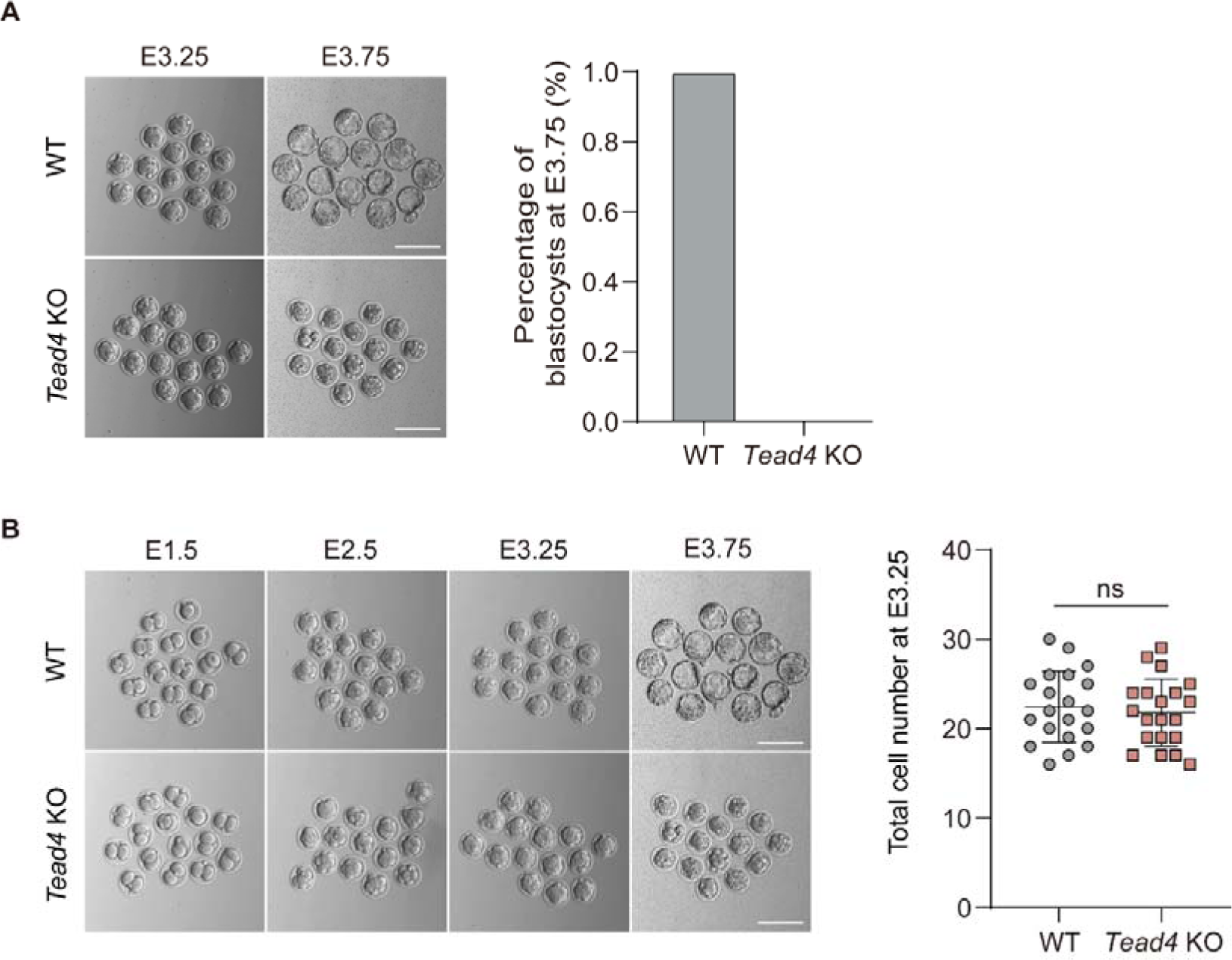
Depletion of *Tead4* leads to developmental arrest at morula stage in mouse. (A) Quantification of the number of blastocysts in control and *Tead4* KO mouse embryos. Scale bar, 25 μm. Data are mean±s.e.m. (n=30). (B) Quantification of total cell number in control and *Tead4* KO embryos at E3.25. Data are mean±s.e.m. (n=40).

### RNA-seq Analysis of Mouse Tead4-deficient Embryos

The developmental phenotypes resulting from disruption of *Tead4* led us to determine the effects of Tead4 on gene expression regulating blastocyst formation at the molecular level. Thus, we collected embryos from WT and KO groups at the late morula stage (E3.25) and then performed RNA-seq (Fig. 3A). Notably, cDNA from each embryo was used for genotyping before library construction (Fig. S2A). Principal component analysis (PCA) showed that two independent replicates of RNA-seq samples from each WT and KO group displayed high correlation (Fig. S2B, C). Moreover, *Tead4* was significantly decreased in the KO group, further confirming robust KO efficiency of *Tead4* by base editor (Table. S2).

**Fig 3.**
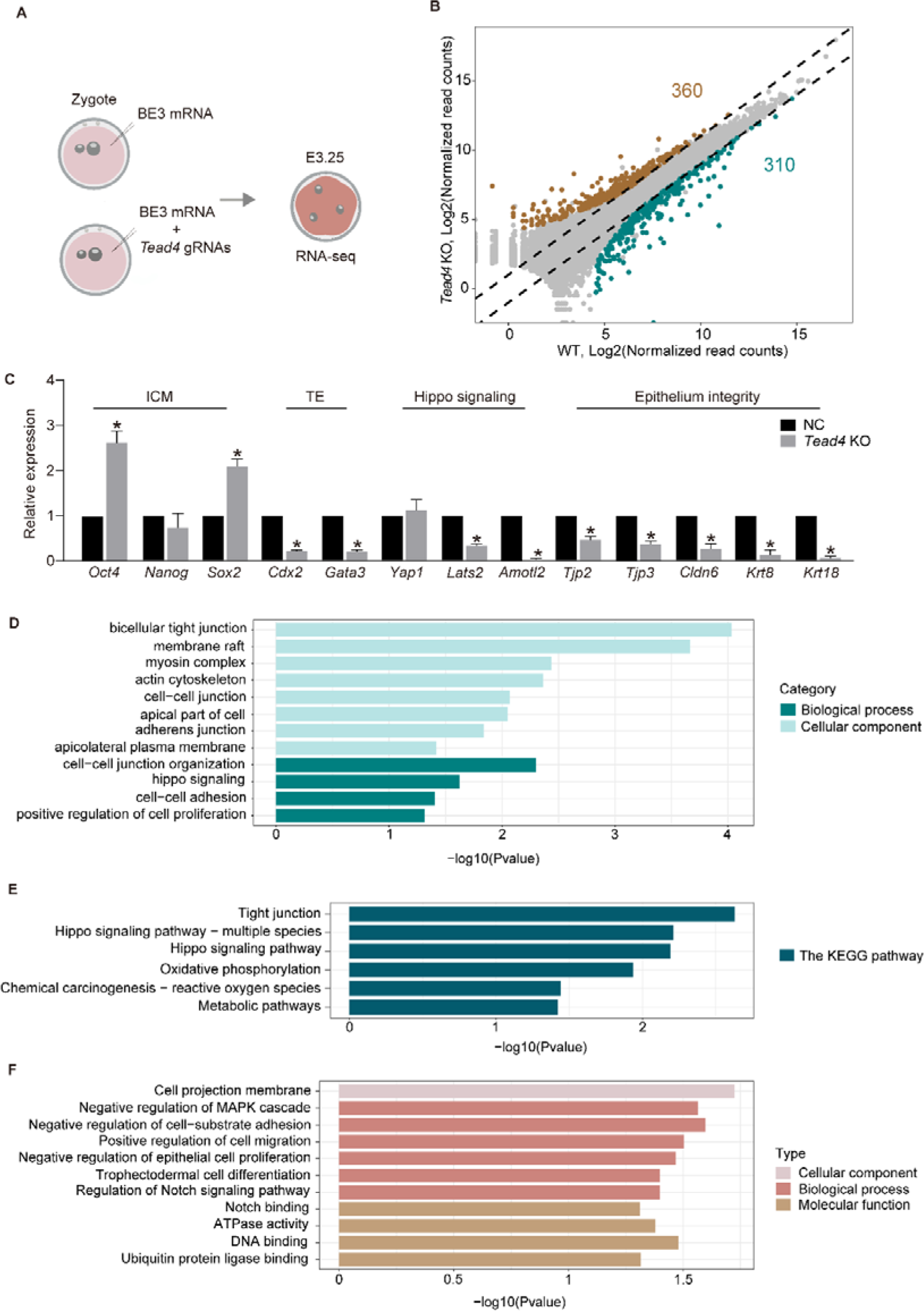
Identification of *Tead4* target genes with RNA-seq in mouse embryo. (A) Scheme of RNA-seq in mouse *Tead4* knockout embryos. Zygotes were injected with BE3 mRNA or *Tead4* gRNAs and BE3 mRNA. Embryos were cultured to morula stage (E3.25) and subjected to genotyping and RNA-seq analysis. (B) Scatter plots showing change of global gene expression in mouse embryos injected with *Tead4* gRNAs and BE3 mRNA compared with embryos injected with only BE3 mRNA at E3.25. (C) Relative expression of genes responsible for ICM specification, TE differentiation, Hippo signaling and epithelium integrity of TE layers (Fold Change >2 or <0.5 and *P*adj <0.05). (D-E) GO (D) and KEGG (E) analysis showing downregulated genes enriched in in tight junction and Hippo signaling pathway (Fold Change >2 or <0.5 and *P*adj <0.05). (F) GO analysis showing upregulated genes enriched in in cell adhesion and epithelial cell proliferation (Fold Change >2 or <0.5 and *P*adj <0.05).

Compared with WT embryos, 670 genes were differentially expressed in *Tead4* KO embryos, with 360 upregulated genes and 310 downregulated genes (Fig. 3B and Table. S2). We observed an increase in the expression of pluripotent genes *Oct4*, *Fgf1*, *Carm1*, *Rhob*, and *Gata4* and a reduction in TE genes *Gata3* and *Cdx2* (Fig. 3C), which aligns with an increased expression of these downregulated genes from morula stage onward in normal embryos (Fig. S2D, E). Additionally, we found a transcriptional repression of tight junction components, such as *Cldn6*, *Tjp3*, *Tjp2*, *Myl6*, *Myl12b*, *Cxadr*, *Cgnl1*, *Clmp*, *Dlg3*, *Amotl2* and *Erbb2*, cytoskeleton component *Krt8* and Hippo signaling components *Lats2* and *Amotl2* compared with WT embryos.

Indeed, gene ontology (GO) enrichment analysis revealed that downregulated genes enriched in cell-cell junctions, tight junctions, apicolateral plasma membrane and Hippo signaling (Fig. 3D), which were similar to the results of Kyoto Encyclopedia of Genes and Genomes (KEGG) pathway analysis (Fig. 3E). Meanwhile, the upregulated genes enriched in negative regulation of cell-substrate adhesion, positive regulation of cell migration and negative regulation of epithelial cell proliferation (Fig. 3F). These data suggest that Tead4 plays functional roles in TE specification, tight junctions and Hippo signaling.

### Tead4 is Required for Mouse Lineage Specification of Trophectoderm

Given the reduction in TE associated markers *Cdx2* and *Gata3* expression through transcriptomic analysis, we asked whether Tead4 could influence TE identity. To address this hypothesis, we performed IF to determine whether there was a reduction at protein level. IF analysis showed a significant decrease in Cdx2 and Gata3 expression compared with WT embryos (Fig. 4A, B). Contrary to Cdx2 and Gata3 expression, disruption of *Tead4* did not alter Sox2 and Nanog expression levels compared with WT embryos (Fig. 4A, C). These results reveal that Tead4 has a significant impact on maintaining TE-specific gene expression but not pluripotency genes in mouse embryos.

**Fig 4.**
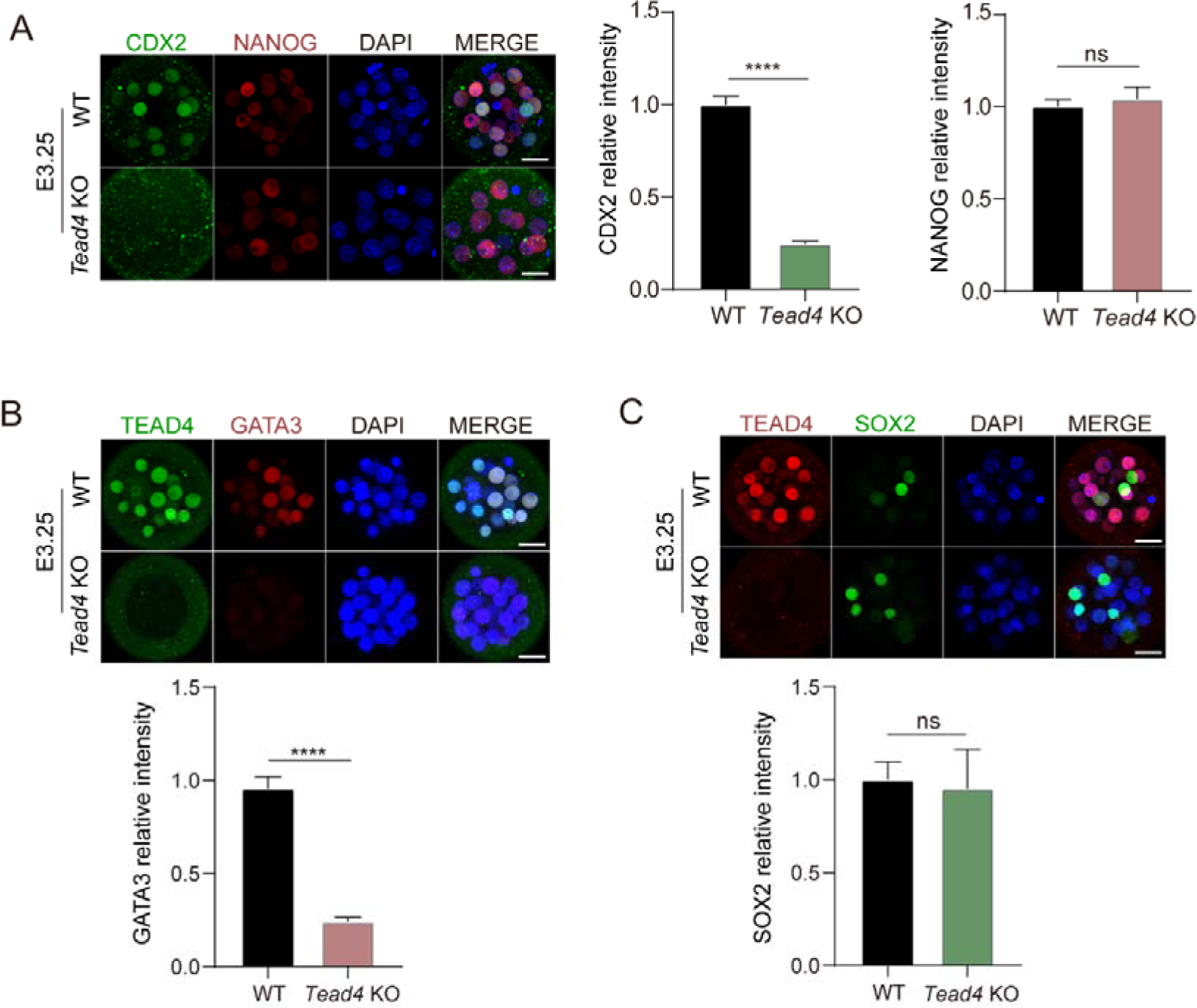
*Tead4* plays a key role in trophectoderm specification in mouse. (A-C) Confocal images and quantification of the relative fluorescence intensity of (A) CDX2 (n=24, 3 biological replicates) and NANOG (n=26), (B) GATA3 (n=19, 3 biological replicates), and (C) SOX2 (n=18, 3 biological replicates) in control and *Tead4* KO mouse embryos at morula stage. Scale bar, 25 μm. Data are mean±s.e.m. ****: P<0.0001.

### Tead4 is Required for Trophectoderm Epithelium Integrity

Epithelium integrity of TE cells is critical for blastocyst formation and expansion ^13^. Our RNA-seq analysis showed a reduction in genes encoding critical components in adhesion junction, tight junction and intermediate components that regulating paracellular integrity. This result raised the question of whether Tead4 may influence TE epithelium integrity in mouse preimplantation embryos. To verify this result, we detected the average intensity of E-Cadherin and β-Catenin (adhesion junction) ^14–16^, TJP2 (tight junction) ^17–19^ and KRT8 (intermediate filament) ^20^. We observed only minor changes in E-Cadherin, β-Catenin and TJP2 (Fig. 5A-C), while there is a significant reduction in KRT8 in *Tead4* KO embryos (Fig. 5D), further substantiating the idea that Tead4 impacts TE epithelium integrity.

**Fig 5.**
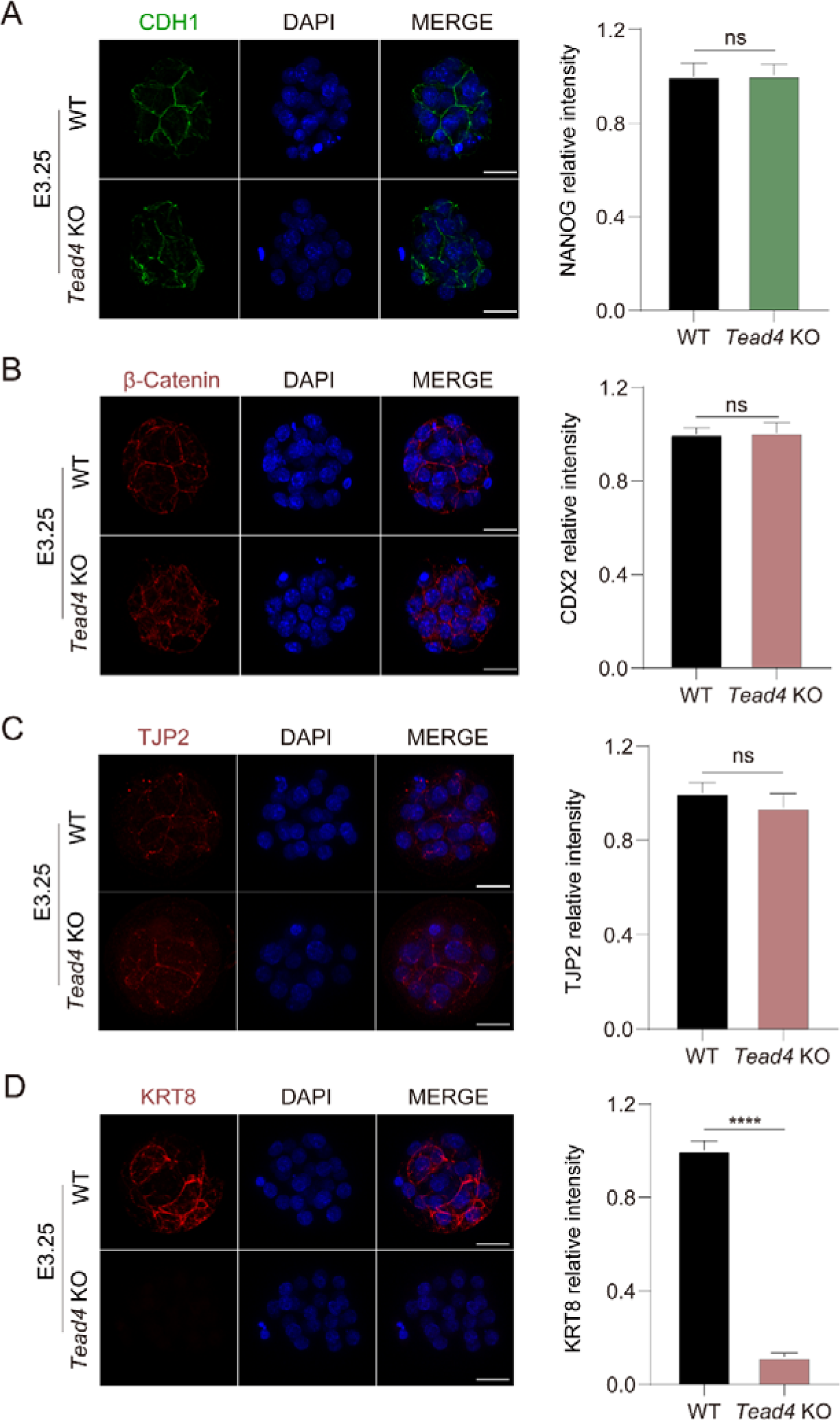
Impaired trophectoderm epithelium integrity in mouse *Tead4* KO embryo. (A-D) Confocal images and quantification of the relative fluorescence intensity of (A) CDH1 (n=26), (B) β-catenin (n=22), (C) TJP2 (n=29, 3 biological replicates) and (D) KRT8 (n=27, 3 biological replicates) in control and *Tead4* KO mouse embryos at morula stage. Scale bar, 25 μm. Data are mean±s.e.m. ****: P<0.0001.

### Tead4 Regulates Hippo Signaling Pathway

RNA-seq analysis also revealed dysregulation of critical components in Hippo signaling pathway, which plays a crucial role in TE lineage specification. As Yap is a core component of Hippo signaling, we examined its expression level and noticed a dramatic decrease in nuclear Yap for *Tead4*-deficient embryos (Fig. 6A), implying that Tead4 directly regulates Hippo signaling in mice.

**Fig 6.**
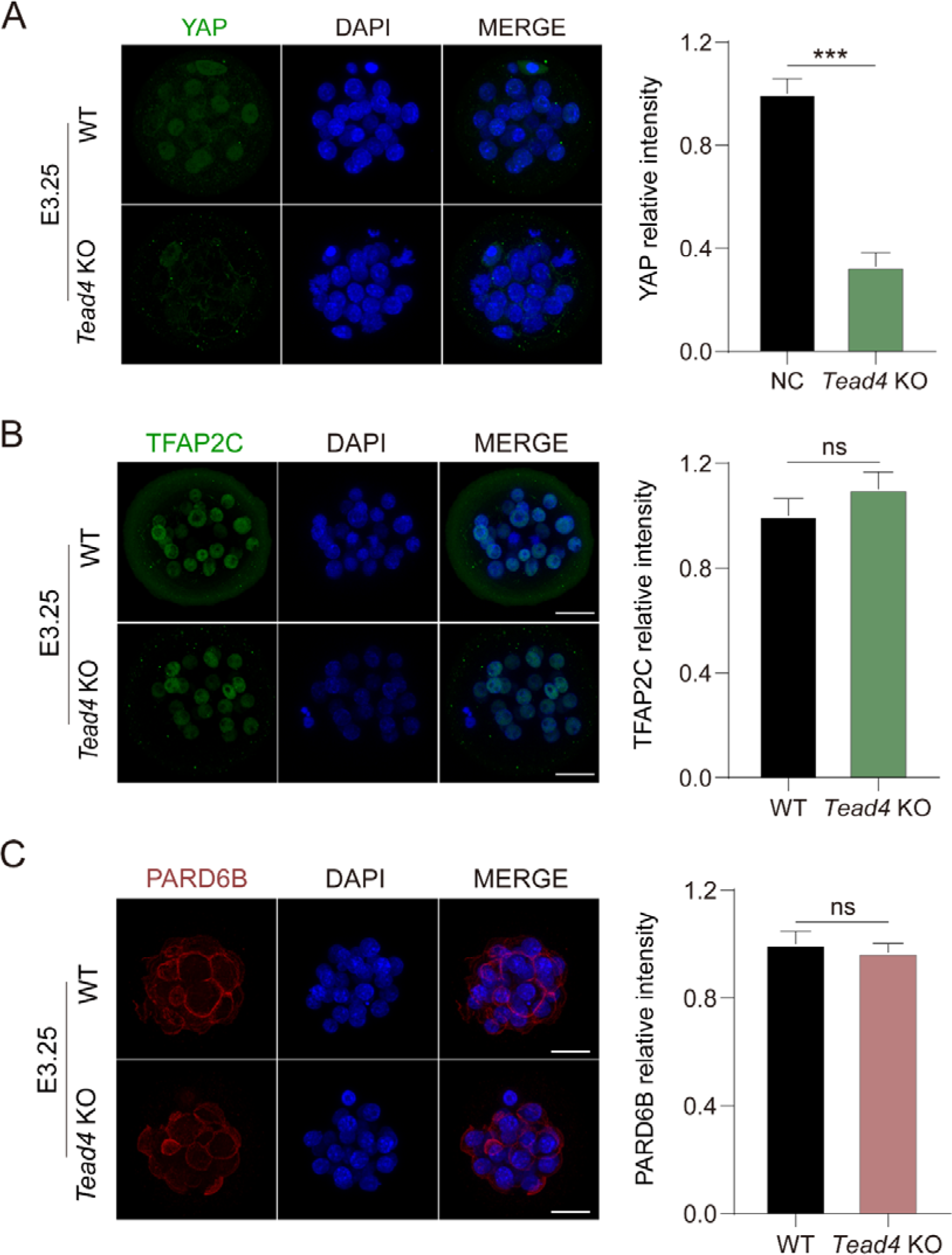
*Tead4* regulates HIPPO signaling not via *Pard6b* in mouse. (A-C) Confocal images and quantification of the relative fluorescence intensity of (A) YAP (n=29, 3 biological replicates), (B) TFAP2C (n=20) and (C) PARD6B (n=14) in control and *Tead4* KO mouse embryos at morula stage. Scale bar, 25 μm. Data are mean±s.e.m. ***: P<0.001.

Moreover, the transcription factor Tfap2c is a key regulator of blastocyst formation and tight junctions ^21^and mirrors the roles of Tead4 we discovered. Furthermore, Tfap2c could regulate the Hippo signaling pathway partly via Pard6b ^22^. Thus, we initially aimed to ascertain whether Tead4 regulates *Tfap2c* expression in mouse embryos. However, we found comparable TFAP2C expression levels between WT and *Tead4* KO groups (Fig. 6B), indicating that Tead4 does not act upstream of Tfap2c in mouse preimplantation embryos. Then, we hypothesized that Tead4 might regulate the Hippo signaling pathway via Pard6b. Interestingly, immunofluorescence analysis revealed a slight change in PARD6B (Fig. 6C). Collectively, our results suggest that Tead4 directly regulates Hippo signaling, but not via Pard6b.

### TEAD4 is Not Required for Blastocyst Formation in Cattle

To investigate if the role of TEAD4 is conserved between mouse and bovine early embryos, we employed BE3 to introduce point mutations in bovine *TEAD4*. We designed two gRNAs in bovine *TEAD4* exon 3 and exon 6 and expected that the premature stop codons would be induced by transition mutation of C4, C5 to T for gRNA1 and C5 to T for gRNA2 at the target DNA sites (Fig. 7A). To evaluate the editing efficiency, we co-injected BE3 mRNA and gRNAs into bovine zygotes (Fig. 7B) and collected the embryos at D8.0 for genotyping. Genotyping results revealed that 85.7% (36/42) of the embryos were correctly edited at target sites for gRNA1, 57.1% (24/42) for gRNA2 and 95.3% (41/43) for gRNA1 or gRNA2 (Fig. 7C). These results demonstrate that base editing provides high editing efficiency of *TEAD4* in bovine embryos.

**Fig 7.**
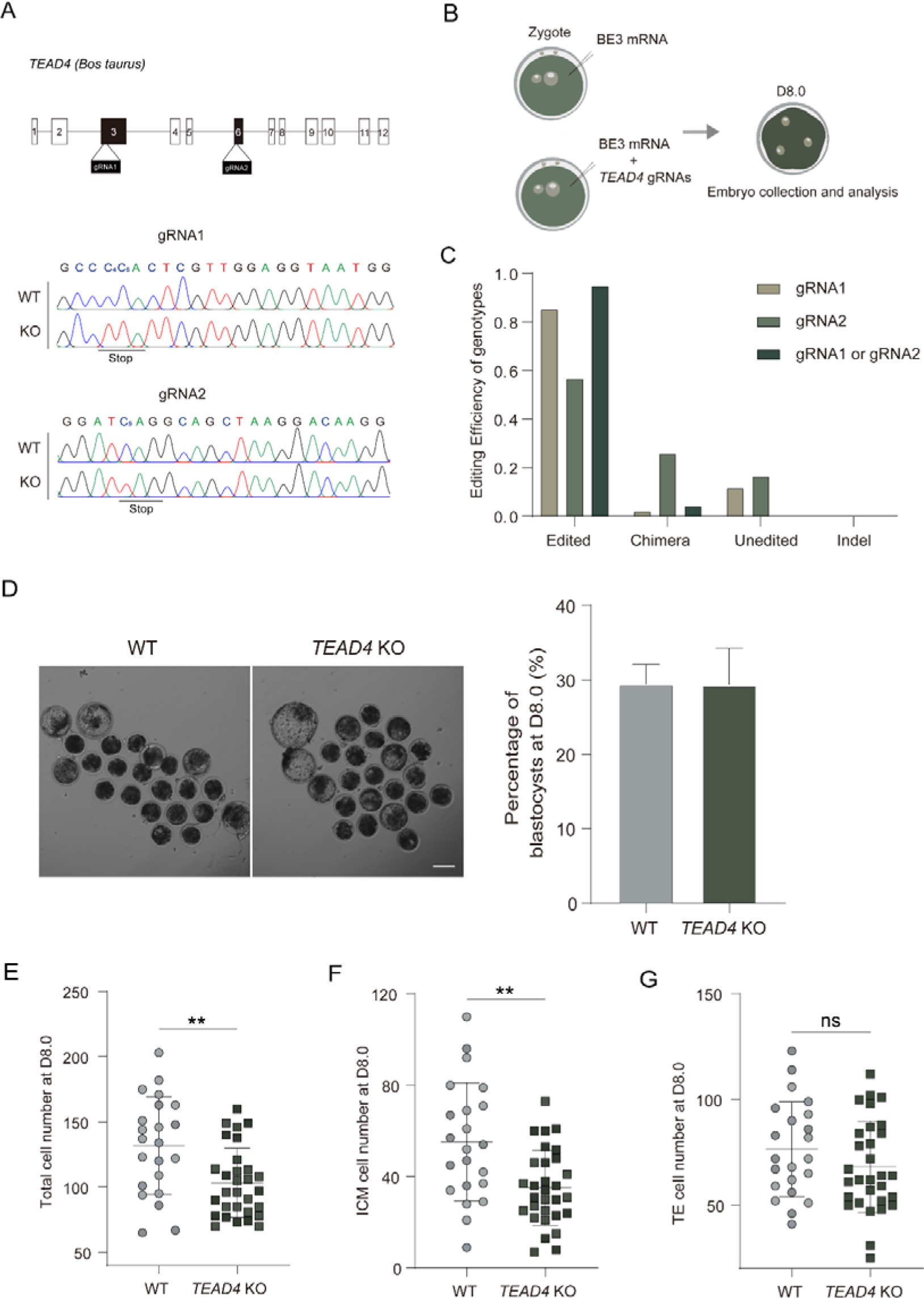
*TEAD4* is not required for blastocyst formation in bovine. (A) Two gRNAs were designed in bovine *TEAD4* exon3 and exon6. C_4_, C_5_ in gRNA1 and C_5_ in gRNA2 are potential target sites. (B) Scheme of bovine *TEAD4* knockout. Zygotes were injected with BE3 mRNA or *TEAD4* gRNAs and BE3 mRNA. Embryos were cultured to blastocyst stage (D8.0) and subjected to genotyping and RNA-seq analysis or immunofluorescence. (C) Editing efficiency of different phenotypes in bovine *TEAD4* KO embryos. Data are mean±s.e.m. (n=43, 4 biological replicates). (D) Quantification of the number of blastocysts in control and *TEAD4* KO bovine embryos. Scale bar, 100 μm. Data are mean±s.e.m. (n=160, 4 biological replicates). (E-G) Quantification of total cell number, ICM cell number and TE cell number in control and *TEAD4* KO bovine embryos. Data are mean±s.e.m. (n=52). **: P<0.01.

We first observed the preimplantation development at D8.0 and found bovine *TEAD4* WT and KO embryos with a comparable number of blastocysts (Fig. 7D). However, the total cell number and ICM cell number of bovine *TEAD4* KO embryos were significantly decreased while the TE cell number was similar (Fig. 7E, F, G), suggesting that cell proliferation of TE was suppressed in *TEAD4* KO embryos. These observations indicate that TEAD4 is not essential for morula to blastocyst transition in bovine embryos.

### TEAD4 is Not Required for Bovine Lineage Specification of Trophectoderm

Despite blastocyst formation in *TEAD4* KO bovine embryos, we sought to gain insight into lineage specification during early development. The staining results indicated that expression of both CDX2 and GATA3, TE associated markers ^7, 23, 24^, were comparable to the WT embryos (Fig. 8A, B, C). Similarly, the expression level of pluripotent gene SOX2 ^1^ barely changed compared with the WT embryos (Fig. 8D, E). Based on the above results, we questioned whether there would be any effects on gene expression in *TEAD4*-depleted embryos before D8.0. We therefore collected embryos at D7.0 and conducted RNA-seq. PCA displayed low correlation between two independent replicates of RNA-seq samples from each WT and KO group (Fig. S4), and the transcriptome data showed only three downregulated genes and no upregulated gene compared with the WT embryos (Table. S3), further supporting the similar developmental capability between two groups. In summary, these data indicate that TEAD4 is not necessary for TE lineage specification in bovine embryos.

**Fig 8.**
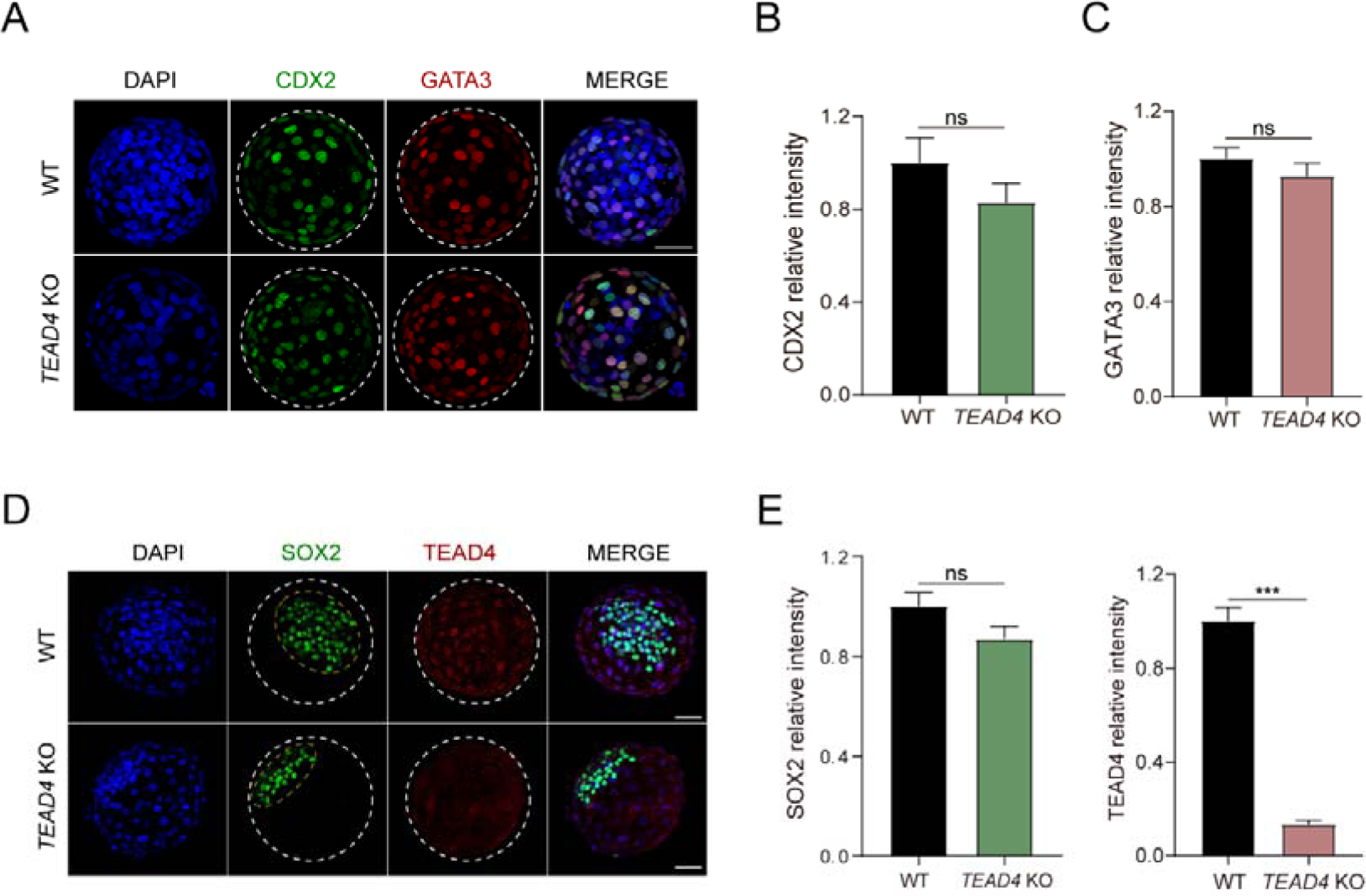
*TEAD4* is not required for trophectoderm specification in bovine. (A-B) Confocal images and quantification of the relative fluorescence intensity of (A-C) CDX2 and GATA3 (n=26), (D-E) SOX2 (n=14, 4 biological replicates) and TEAD4 (n=42, 3 biological replicates) in control and *TEAD4* KO mouse embryos at blstocyst stage (D8.0). Scale bar, 50 μm. Data are mean±s.e.m. ***: P<0.001

## Discussion

How the first lineage specification event in mammals is resolved is a fundamental question in developmental biology. This study explores the potential mechanisms underlying the functional requirement of Tead4 in the first lineage specification in mice and determines the dispensable role of TEAD4 in bovine preimplantation embryos. Three novel findings have emerged from this study. Firstly, we have identified a dysregulation of 670 genes in *Tead4* KO embryos. Secondly, we have observed a reduction in KRT8, suggesting that Tead4 influences the integrity of the TE epithelium in mice, and a decrease in nuclear YAP for Tead4-deficient embryos, indicating that Tead4 directly regulates Hippo signaling. Lastly, we have found that bovine embryos with TEAD4 depletion can still develop to blastocysts with normal expression of *CDX2*, *GATA3*, and *SOX2*. In summary, we propose that Tead4 regulates mouse blastocyst formation via Krt8 and Yap, both of which are critical regulators of mouse preimplantation development (Fig. 9).

**Fig 9.**
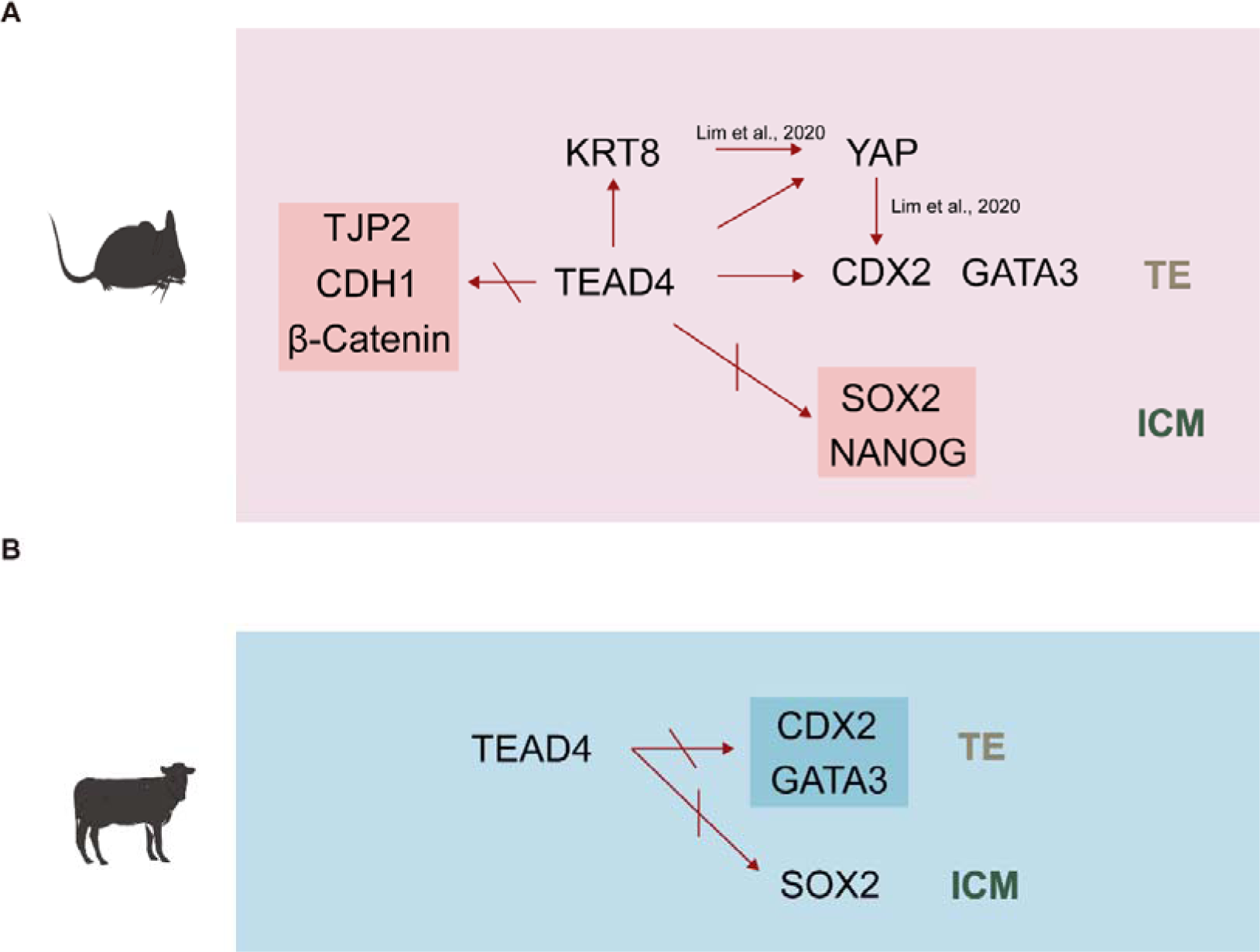
Model of TEAD4 transcriptional regulation in mouse and bovine embryo. For mouse, TEAD4 is crucial for trophectoderm specification and epithelium integrity. For bovine, TEAD4 is not required for lineage specification.

We established that Tead4 is required for blastocyst formation and TE lineage specification by using a base editing approach to induce *Tead4* ablation. This is consistent with earlier studies using homologous recombination. Recently, scientists also used WT Cas9 to study gene function in mammalian preimplantation embryos. However, the genome editing with WT Cas9 exhibits low editing efficiency and induces mosaic genotypes, which complicates the phenotypic analysis and developmental assessment of the injected embryos. Base editors, derived from CRISPR/Cas9 mediated genome editing, can be used to precisely install target point mutations with fewer undesired byproducts and without double-strand DNA breaks (DSBs), donor DNA templates and homology directed repair (HDR) ^25, 26^. In this study, our two gRNAs exhibited about 80% targeted editing and injection of the mixture resulted in almost 100% targeted editing efficiency in the embryo we used for RNA-seq and IF. Thus, base editing represents a robust and reliable approach to dissect gene function in mammalian early embryos, especially helpful for animals for which germline genetic engineering is challenging.

It is believed that polarity-Hippo/Yap signaling plays a key role in TE initiation in mammals. The Hippo signaling pathway acquires different states in inner and outer cells of the embryo, resulting in different fates. Unphosphorylated Yap/TAZ enters the nucleus of outer cells, binds to Tead4, and activates the expression of TE lineage specifiers. Interestingly, *Tead4* KO results in dysregulation of several genes encoding key components of the Hippo signaling pathway, including *Lats2* and *Amotl2*. Importantly, the signal intensity of Yap in the nuclear region is drastically reduced after *Tead4* ablation, indicating a failure of Yap to translocate to outer cells and eventually triggering a failed TE specification.

Krt8 and Krt18 are key components of intermediate filaments, enriched at the apical domain and inherited asymmetrically by outer cells ^20^. Krt8 accomplishes translocated expression from nuclear to cell borders and expresses exclusively in the TE cell layer but not inner cell mass at late blastocyst stage in mouse preimplantation embryos ^27, 28^, indicating Krt8 as a marker of TE in mouse. Krt8 ablation results in a decrease in Yap, a transcriptional binding factor and Cdx2 exhibited reduced expression in Yap-depleted embryos in mice ^20^, reflecting that Krt8 regulates Cdx2 through Yap in mouse preimplantation embryos. We then found a reduction in Krt8 and Cdx2 in *Tead4* disrupted mouse embryos (Fig. 4A; 5D). Taken together, Tead4 could regulate Cdx2 directly or through Krt8-Yap (Fig. 9A).

Recently, two studies showed that knocking down *TEAD4* via RNA interference does not affect development to the blastocyst stage in cattle, but they produced contrasting results regarding the role of TEAD4 in TE marker genes. Specifically, Hiroki Akizawa et al. found that knocking down *TEAD4* significantly reduced the transcript levels of *CDX2* but did not affect the expression of *GATA3* in bovine embryos ^29^; while Nobuyuki Sakurai et al. found that *TEAD4*, *CDX2*, *GATA3*, *OCT4* and *NANOG* transcripts were not affected in Tead4 knockdown bovine embryos ^30^. To resolve this issue, our data clearly show that *TEAD4* KO does not affect TE and ICM lineage markers. Considering highly conserved TEAD family proteins, it is possible that other TEADs could play a compensatory role with TEAD4, which warrants further study.

In summary, we confirm the essential role of Tead4 in blastocyst formation and TE specification in mouse preimplantation embryos. Our data suggest this functional role is likely mediated by Krt8 and Yap. Moreover, Tead4 is dispensable for blastocyst formation and TE lineage specification in cattle.

## Materials and methods

### Ethics statement

Animals were maintained in accordance with the Guidelines for Ethical Review of laboratory Animal Welfare and approved by Zhejiang University.

### Mouse embryo collection and *in vitro* culture

Embryos were collected from 8-10-week-old B6D2F1 (C57BL/6×DBA/2; Beijing Vital River Laboratory Animal Technology) superovulated female mice with 8 international units (IU) of pregnant mares’ serum gonadotropin (PMSG; Sansheng Pharmaceutical) and 8 IU human chorionic gonadotropin (hCG; Sansheng Pharmaceutical) 46-48 hours later, and then crossed with B6D2F1 male mice. Embryos were isolated from oviducts in M2 medium (MilliporeSigma) and transferred to hyaluronidase solution (MilliporeSigma) to remove cumulus cells and cultured in KSOM medium (MilliporeSigma) at 37[with 5% CO_2_. Mouse embryos were fixed at the following times post fertalization: one-cell stage (12h post fertilization, E0.5), two-cell stage (36h post fertilization, E1.5), four-cell stage (48h post fertilization, E2.0), eight-cell stage (60h post fertilization, E2.5), sixteen-cell stage (72h post fertilization, E3.0), morula stage (78h post fertilization, E3.25), early blastocyst stage (90h post fertilization, E3.75), late blastocyst stage (102h post fertilization, E4.25).

### Bovine embryo *in vitro* production

Procedures for bovine embryo *in vitro* production includes *in vitro* maturation (IVM), *in vitro* fertilization (IVF) and *in vitro* culture (IVC). Cumulus-oocyte complexes (COCs) with more than three cumulus cell layers were cultured in IVM medium at 38.5[with 5% CO_2_ for 22-24 h. The IVM medium contains Medium-199 (Sigma), 10% fetal bovine serum (FBS) (Gibco), 1 IU/ml follicle-stimulating hormone (FSH) (Sansheng Biological Technology), 0.1 IU/ml luteinizing hormone (LH) (Solarbio), 1 mM sodium pyruvate (Thermo Fisher Scientific), 2.5 mM GlutaMAX (Thermo Fisher Scientific) and 10 μg/ml gentamicin. Matured COCs were incubated with spermatozoa (1-5×10^6^/ml) purified from thawed semen with Percoll in BO-IVF medium (IVF Bioscience) at 38.5[with 5% CO_2_ for 9-12 h. The cumulus cells were removed with 1 mg/ml hyaluronidase, followed by embryo culture in BO-IVC Medium (IVF Bioscience) until late blastocyst stage (192 h post fertilization, D8.0).

### Creation of zygotic Tead4 knockout

Guide RNAs (gRNAs) were designed on BE-designer (http://www.rgenome.net) and synthesized with 5’ extended CACC and AAAC (Sangon Biotech) on the two DNA strands respectively before annealed to double strands. The DNA oligos were then ligated to PX458 vector linearized by BpiI (Takara) and cloned into *E.coli* DH5α competent cells (Takara). Plasmids were extracted from bacterial cultured overnight, followed by amplification as polymerase chain reaction (PCR) templates. The PCR primers contain T7 promoter. The sequences of gRNAs and primers are listed in Table S4.

### *In vitro* transcription and microinjection

BE3 plasmid was purchased from Addgene (Cat# 73021) and linearized with NotI, followed by purification with GeneJET PCR Purification Kit (Thermo Fisher Scientific) and in vitro transcription with mMESSAGE mMACHINE T7 Ultra Kit (Thermo Fisher Scientific). gRNAs were in vitro transcribed with MEGAshortscript T7 High Yield Transcription Kit (Thermo Fisher Scientific). Controls were injected with BE3 mRNA (final concentration, 200 ng/μl) and KOs were injected with a combination of two gRNAs (final concentration, 100 ng/μl respectively) and BE3 mRNA (final concentration, 200 ng/μl). mRNAs were microinjected into cytoplasm of zygotes 20-22 h post hCG injection, using an Eppendorf transferman micromanipulators.

### Genotyping of single embryo

Each embryo was individually collected into a tube containing lysis buffer (40 nM Tris-HCl, 1% NP-40, 1% Triton X-100 and 0.4 ng/ml Proteinase K) and then incubated at 55[for 1 h and 95[for 10 min as PCR template. Nested PCR was performed with rTaq (Takara), followed by the PCR product fragments sequencing with Sanger sequencing. The sequences of primers are listed in Table S4 and the Nested PCR reaction conditions are listed in Table S5.

### Immunofluorescence

Embryos were washed third in 0.1% PBS/PVP (PBS containing 0.1% PVP) and fixed in 4% paraformaldehyde (PFA) for 10 min (bovine, 30 min) at room temperature (RT), followed by permeabilization in PBST (PBS containing 0.5% Triton X-100) for 30 min (bovine, 40 min). Embryos were then blocked in 10% FBS in 0.1% Triton X-100/PBS for 1 h (bovine, 2 h) and incubated with primary antibodies in blocking solution overnight at 4[. The embryos were washed third in 0.1% Triton X-100/PBS and incubated with secondary antibodies in blocking solution for 1 h (bovine, 2 h) before washed in 0.1% Triton X-100/PBS. The embryos were then incubated with DAPI in PBS for 10 min (bovine, 20 min) and imaged in drops of 0.1% Triton X-100/PBS on glass slide with Zeiss LSM 880 confocal microscope. All antibodies involved in this research are listed in Table S6.

### RNA-seq library construction and sequencing

Mouse embryos were collected after removal of the zona pellucida with Tyrode’s solution at morula stage (n=24, 2 biological replicates). Total RNA was extracted with Arcturus PicoPure RNA Isolation Kit (Life Technologies). Then mRNAs were fragmented using oligo(dT)25 beads and reverse transcribed. Sequencing libraries were constructed with NEB Next Ultra RNA Library Prep Kit for Illumina (New England Biolabs). The cDNA was preamplified with KAPA HiFi HotStart ReadyMix, purified with Ampure XP beads, fragmented by Tn5 enzyme (Vazyme) and amplified for 15-18 cycles before paired-end 150 bp sequencing on Illumina NovaSeq (Novogene).

### RNA-seq data alignment and analysis

The raw sequencing reads were trimmed with Trimmomatic (version 0.39) ^31^ to remove adaptor sequences and low quality reads and obtain clean reads which were then mapped to mm10 with Hisat2 (version 2.1.0) ^32^. The raw read counts were calculated with featureCounts (version 1.6.3) ^33^ and then normalized to FPKM with Cufflinks (version 2.2.1) ^34^. The differentially expressed genes were identified by DESeq2 with fold change >2 or <0.5 and adjusted *P*-value <0.05. Enrichment analysis of differentially expressed genes was performed with the Database for Annotation, Visualization and Integrated Discovery (DAVID) ^35, 36^.

### Statistical analysis

Charts and statistics were generated in GraphPad Prism 8.0 and ImageJ. Quantitative analysis was performed with two-tailed unpaired Student’ s t-test and is presented as mean±standard error of the mean (s.e.m.). P<0.05 indicates significant difference.

## Acknowledgements

We are grateful to members of K.Z. lab for their helpful discussions and comments on this manuscript. We thank laboratory of animal center of Zhejiang University for taking care of mice, analysis center of agrobiology and environmental sciences of Zhejiang University for Zeiss LSM 880.

## Author contributions

Conceptualization, X.W., Y.S. and K.Z.; Investigation, X.W., Y.S., P.Z., B.H., S.L. and L.X.; Formal analysis, X.W., Y.S., B.H. and S.L.; Writing - original draft, X.W.; Writing - review & editing, S.W. and K.Z.; Funding acquisition, K.Z.; Supervision: S.W. and K.Z.

